# Mountain radiations are not only rapid and recent: Ancient diversification of South American frog and lizard families related to Paleogene Andean orogeny and Cenozoic climate variations

**DOI:** 10.1101/2021.04.24.441240

**Authors:** Lydian M. Boschman, Fabien L. Condamine

## Abstract

Mountainous areas host a disproportionately large fraction of Earth’s biodiversity, suggesting a causal relationship between mountain building and biological diversification. Mountain clade radiations are generally associated with changes in environment, climate, and the increase in heterogeneity therein during mountain building. However, examining the causal relationship between mountain building and diversification is a complex challenge, because isolating the effects of surface uplift from other abiotic (climate) or biotic variables is not straightforward. In this study, we investigate the relative contributions of abiotic climate-driven (temperature) and geology-driven (elevation) drivers on evolutionary rates of ancient groups of organisms in the South American Andes. We present regional curves of Andean elevation based on a recent compilation of paleo-elevational data back to the Late Cretaceous, and analyse the diversification history of six Andean frog and lizard families that originated equally far back in time. For two clades (Aromobatidae and Leptodactylidae), we find that they diversified most rapidly during the early phase of mountain building (Late Cretaceous - Paleogene), when the first high-elevation habitats emerged in South America. The diversification of one clade (Centrolenidae) is correlated with Cenozoic temperature variations, with higher speciation rates during warm periods. The last three clades (Dendrobatidae, Hemiphractidae and Liolaemidae) are best explained by environment-independent diversification, although for Liolaemidae, an almost equally strong positive correlation was found between speciation and Andean elevation since the late Eocene. Our findings imply that throughout the long-lived history of surface uplift in the Andes, mountain building drove the diversification of different clades at different times, while not directly affecting other clades. Our study illustrates the importance of paleogeographic reconstructions that capture the complexity and heterogeneity of mountain building in our understanding of the effects that a changing environment plays in shaping biodiversity patterns observed today.

**Highlights:**

- We provide novel regional paleoelevation curves for the Andes back to the Late Cretaceous
- The diversification history of six Andean-centered clades is studied
- We find clade-specific responses to environmental changes
- The impact of Andean uplift could reach further back in time than previously thought

## 1. Introduction

Mountainous areas cover approximately 25% of the surface of the continents, yet are home to 85% of the world’s terrestrial amphibian, bird, and mammal species (Körner et al. 2016; Rahbek et al. 2019b). This strong imbalance in species richness suggests a causal relationship between mountain building and species radiations (Antonelli et al. 2018a; Hoorn et al. 2018; Rahbek et al. 2019a), and such radiations have been documented for various mountainous regions, across disparate groups of organisms (e.g. Drummond et al. 2012; Schwery et al. 2015; Favre et al. 2016; Lagomarsino et al. 2016; Ebersbach et al. 2017; Hutter et al. 2017; Xing and Ree 2017; Condamine et al. 2018; Esquerré et al. 2019; Muellner-Riehl et al. 2019; Ye et al. 2019; Ding et al. 2020). Mountain clade radiations are generally explained by landscape changes and the associated changes in climate and habitat connectivity, which lead to ecological opportunities and evolutionary innovations (Drummond et al. 2012; Hughes and Atchinson 2015; Favre et al. 2016; Lagomarsino et al. 2016; Cortés et al. 2018). As a result, mountain clade radiations can be expected to be coeval with the timing of surface uplift. Although geological evidence for uplift histories of mountain ranges come with large uncertainties (Blisniuk and Stern 2005; Rowley and Garzione 2007; Botsyun et al. 2020) and uplift is often spatially heterogeneous (e.g. Liu et al. 2016; Spicer et al. 2020; Boschman 2021), most of the present-day mountain habitats across the globe are thought to have formed gradually throughout the Cenozoic (e.g. Chamberlain et al. 1999, 2012; Nakajima et al. 2006; Fauquette et al. 2015; Liu et al. 2016; Boschman 2021). It is therefore surprising that documented mountain radiations are found to be generally recent (largely confined to the Pliocene and Pleistocene) and rapid (Linder 2008; Hughes and Atchinson 2015; Quintero and Jetz 2018).

The South American Andes, stretching over ∼7,000 km from tropical Colombia to sub-polar Patagonia, are the most biodiverse mountains in the world, and the northern, Tropical Andes are considered the most species-rich of all biodiversity hotspots (Myers et al. 2000; Mittermeier et al. 2004). Including some of the most diverse radiations on Earth (Hughes and Eastwood 2006; Antonelli and Sanmartín 2011; Madriñan et al. 2013; Lagomarsino et al. 2016; Pérez-Escobar et al. 2017; Esquerré et al. 2019; Testo et al. 2019), this exceptional species richness is thought to be primarily the result of environmental heterogeneity along both the latitudinal and altitudinal gradients, resulting in a wide variety of climates, landscapes and vegetation types including dry forests and woodlands, tropical rainforests, cloud forests, and permanently or seasonally snow-covered grasslands (Squeo et al. 1993; Josse et al. 2011; Luebert and Weigend 2014; Cuesta et al. 2017). However, in addition, the paleoclimatic and paleogeographic history of the South American continent and of the Andes in particular is thought to have played a significant role (Gentry 1982; Antonelli and Sanmartín 2011). Since the break-up of the supercontinent Pangea and Late Cretaceous separation from Africa, South America remained an isolated “island-continent” (Simpson 1980) until the late Miocene formation of the Panama Isthmus (Montes et al. 2015; O’Dea et al. 2016), and shifted little in latitude. These ∼90 Myr of isolation and relative climatic stability may have favoured endemism and the gradual accumulation and preservation of lineages (Antonelli and Sanmartín 2011). Nonetheless, global-scale climate fluctuations and mountain building led to significant variations in environment (Armijo et al. 2015), affecting the evolution of life. For example, plant diversity has been shown to be positively correlated with Cenozoic temperature, whereby diversity levels during the warm Eocene likely exceeded the Holocene and present (Wilf et al. 2003; Jaramillo et al. 2006). Andean mountain building affected diversity in multiple ways: locally, mountain building is thought to have increased diversity through isolation and allopatric speciation (Hazzi et al. 2018) and ecological adaptation to altitude (Nevado et al. 2018). Regionally, the Andes are often considered to have acted as a “species pump”, producing lineages that colonized the surrounding Neotropical lowlands (e.g. Santos et al. 2009; Luebert and Weigend 2014; Chazot et al. 2019). Moreover, the environmentally diverse slopes of the Andes have repeatedly attracted non-Andean lineages, acting as a “species attractor”, increasing species richness through colonization (e.g. Drummond et al. 2012; Chazot et al. 2016; Hutter et al. 2017; Toussaint et al. 2019). These mechanisms are not mutually exclusive because for some clades, a mixture of both processes (colonization in and out of the Andes) has been reported, increasing diversity in both the mountains and the surrounding lowlands (Brumfield and Edwards 2007; Pérez-Escobar et al. 2017; Antonelli et al. 2018b).

The many recent (Neogene) documented mountain radiations in and around the Andes are commonly explained in light of recent uplift (Hughes and Eastwood 2006; Hoorn et al. 2010; Antonelli and Sanmartín 2011; Madriñan et al. 2013; Luebert and Weigend 2014; Lagomarsino et al. 2016; Pérez-Escobar et al. 2017; Esquerré et al. 2019; Testo et al. 2019). This recent uplift is inferred from studies presenting stable isotope paleoaltimetry and fossil leaf physiognomy data from the Eastern Cordillera of Colombia and the eastern Altiplano/Eastern Cordillera of the central Andes, concluding that rapid uplift occurred in the last ∼12 million years (Gregory-Wodzicki 1998, 2000; Garzione et al. 2008). However, the sedimentary records from the Andean foreland basins indicate that uplift in the Andes initiated already in the Late Cretaceous: ∼100 million years ago (Ma) in Patagonia, and ∼70 Ma in the central and northern Andes (Horton 2018). Moreover, since the studies of Gregory-Wodzicki (1998; 2000) and Garzione et al. (2008), a wealth of paleoaltimetry datasets has been published, depicting a much more complex picture (e.g. Bershaw et al. 2010; Leier et al. 2013; Canavan et al. 2014; Carrapa et al. 2014; Garzione et al. 2014; Hoke et al. 2014; Saylor and Horton 2014; Quade et al. 2015; Anderson et al. 2015, 2016; Fiorella et al. 2015; Kar et al. 2016; Rohrmann et al. 2016; Colwyn et al. 2019). This body of work, compiled and summarized in Boschman (2021), shows that (i) topography was already in place in Patagonia and in the western ranges along the Pacific coast during the Late Cretaceous and Paleogene, (ii) most of the central and eastern ranges were uplifted during the last 50 to 30 Myr, and (iii) uplift migrated further eastwards towards the Subandean zone in the last 10 Myr (**Fig. 1**). Nonetheless, the idea that the bulk of the topography in the Andes has emerged very recently has persisted in biogeographic and macroevolutionary literature (e.g. Hughes and Eastwood 2006; Lagomarsino et al. 2016; Pérez-Escobar et al. 2017; Hazzi et al. 2018).

**Figure 1.**
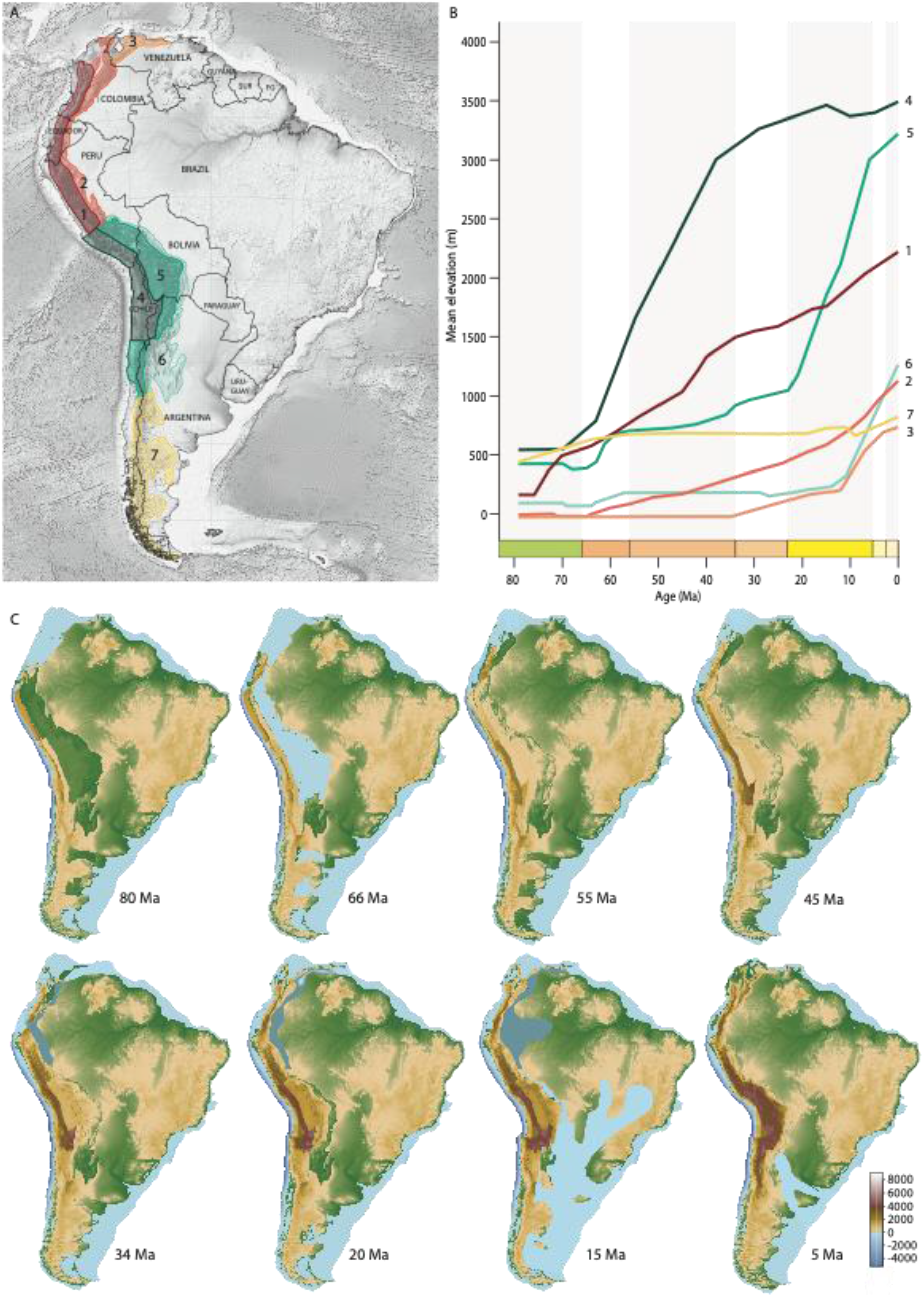
A) Map of South America; digital elevation model in black and white from Etopo1 (Amante and Eakins 2009). The Andes are divided into seven geomorphological domains, based on uplift history. B) Paleoelevation history of the seven domains since 80 Ma, showing the spatial and temporal heterogeneity of the uplift. Negative average elevations (of northern Andean domains) represent elevations below sea level. C) Maps of Andean mountain building. The maps include marine incursions (in light blue) and lakes and wetlands (in grey-blue) based on Hoorn and Wesselingh (2001) for Amazonia and Hernández et al. (2005) for southeastern South America. Topography of the eastern part of the continent is not reconstructed (modern topography is shown) and the maps do therefore not include much less well-documented changes in topography in the Guiana and Brazilian shields areas, or other paleogeographic features. Paleoelevation curves and maps are based on Boschman (2021).

Examining the causal relationship between mountain building and diversification is a complex challenge, because isolating the effects of surface uplift throughout the Cenozoic from other abiotic (climate) or biotic variables is not straightforward (Marx and Uhen 2010; Ezard et al. 2011, 2016; Condamine et al. 2019a,b). Moreover, these abiotic and biotic variables are not independent, as mountain building affects both local and regional climate and biotic mechanisms and interactions, which, in turn, have their effects on diversification (Hoorn et al. 2010; Favre et al. 2015; Antonelli et al. 2018; Ding et al. 2020). Because of this synergy of factors that can affect diversification, there is debate over whether mountain uplift was the primary factor in promoting diversification or that instead, climate change may have played a crucial role (Drummond et al. 2012; Hoorn et al. 2013; Mutke et al. 2014; Hughes and Atchison 2015; Hutter et al. 2017; Nevado et al. 2018). An important step towards answering such questions is the quantitative correlation between diversification rates and environmental variables. Four studies so far, focussing on radiations of young (Neogene) clades, have attempted this, and found positive correlations between speciation rates and the paleoelevation of the Andes, suggesting a causal link between surface uplift and diversification of the studied clades (Lagomarsino et al. 2016; Pérez-Escobar et al. 2017; Esquerré et al. 2019; Testo et al. 2019). However, these studies have applied elevation-dependent diversification models, without comparing, in a common statistical framework, other diversification models considering the effect of time alone, or of other environmental variables (e.g. temperature). In this study, we embrace the challenge to study the relative contributions of climate-driven abiotic and geology-driven abiotic drivers on evolutionary rates of ancient groups of organisms. We aim at teasing apart the contribution of Andean mountain building, for which we use a novel reconstruction that includes surface uplift since the Late Cretaceous, global climate change, and time, on the diversification of Neotropical frog and lizard groups mostly distributed in the Andes. This study provides the next step towards an understanding of why some lineages diversify extensively during uplift while others do not, and consequently, of our understanding of modern biodiversity patterns including the relationship between topography and species richness.

## 2. Materials and methods

### 2.1. Selection of biological groups occurring in the Andes

We focus in this study on amphibians and squamates that are particularly well diversified in the Andes (Myers et al. 2000; Mittermeier et al. 2004). We compiled a dataset of species-level time-calibrated phylogenies from the literature, including family-level phylogenies with a rich species diversity that sampled at least 65% of the total species diversity (i.e. sampling fraction of 0.65: ratio of sampled species over known species). For amphibians, we relied on a phylogeny of 2,318 hyloid frog species (Hutter et al. 2017), which was constructed with a supermatrix analysis of molecular data. There are 2,488 known species of Hyloidea in South America, and 1,594 were sampled in the tree (64%). From this tree, we selected the five best sampled frog families with an average sampling fraction of 0.793 (654 sampled species over 825 known species) and with an average age of origin of 65.4 Ma (**Table 1**). For squamates, we only selected the lizard family Liolaemidae, which is one of the richest vertebrate Andean radiation with over 320 species (258 sampled), originating around 37 Ma (**Table 1**). The species-level time-calibrated phylogeny was retrieved from the study of Esquerré et al. (2019). Four of the six selected clades originated during the Late Cretaceous; two are younger and originated during the late Eocene (Liolaemidae) or at the Eocene-Oligocene boundary (Centrolenidae).

**Table 1.** Summary of diversity, distribution, and ages of hyloid frog and lizard clades used to estimate the diversification rates. The frog data are from the study of Hutter et al. (2017) with updates from the AmphibiaWeb (http://amphibiaweb.org/), while the lizard data are from the study of Esquerré et al. (2019) with updates from the Reptile Database (http://www.reptile-database.org/).

| Clade | Age of origin (Ma) | Number of genera | Species diversity (sampled) | Sampling fraction | Elevational range (m) | Distribution range |
| --- | --- | --- | --- | --- | --- | --- |
| Aromobatidae | 67.18 | 5 | 127 (118) | 0.929 | 500 (0-3300) | Northern and central Andes, Amazonia, Atlantic Forest |
| Centrolenidae | 33.4 | 12 | 160 (128) | 0.8 | 1400 (0-3501) | Northern and central Andes, Amazonia, Atlantic Forest |
| Dendrobatidae | 67.35 | 16 | 197 (136) | 0.69 | 840 (0-3799) | Northern and central Andes, Amazonia |
| Hemiphractidae | 80.69 | 6 | 118 (86) | 0.729 | 1640 (0-4600) | Northern and central<br>Andes, Atlantic<br>Forest |
| Leptodactylidae | 78.07 | 14 | 223 (186) | 0.834 | 500 (0-4480) | whole South America |
| Liolaemidae | 36.9 | 3 | 321 (258) | 0.804 | 2150 (20-5000) | Central and southern<br>Andes |

The six selected clades are widespread in either the whole (Leptodactylidae), central and southern (Liolaemidae), or northern and north-central Andes (the other four; **Supplementary materials**). However, except for Liolaemidae, they are not confined to the Andean ranges alone, and for Aromobatidae and Leptodactylidae, the main center of species diversity is in fact outside of the Andes: in Amazonia and the Guiana Shield region, and in the Atlantic Forest, respectively. Species of the genus *Adenomera* are for example most widely distributed throughout the lowlands of tropical South America east of the Andes (Fouquet et al. 2014). Nonetheless, the origin and geographic range evolution of Aromobatidae and Leptodactylidae are thought to be intimately linked to the Andes (Santos et al. 2009; Hutter et al. 2017), and most species in these clades Aromobatidae and Leptodactylidae are associated with montane environments and are leaf-litter dwelling, which favors allopatry (Fouquet et al. 2013; Santos et al. 2020).

### 2.2. Reconstruction of Andean uplift

We used the reconstruction of paleoelevation in the Andes since 80 Ma of Boschman (2021), which is based on a wide variety of input data, including stable isotope paleoaltimetry, stratigraphy, thermochronology, paleosurfaces, paleobotany, paleontology, palynology and fossil leaf physiognomy. This reconstruction is presented as a series of raster files, one per million-year time step, in 0.1 degree resolution. It thereby provides a very detailed overview of the history of Andean mountain building, including the stark contrast in timing and magnitude of uplift between the different domains of the Andes. To convert this reconstruction (**Fig. 1C**) into a quantitative time series that we can use in the birth-death modelling, we first condensed it into uplift curves (in m.a.s.l. through geological time) for seven individual geomorphological domains (**Fig. 1A, B**) for which we describe the history of uplift below. Second, we computed curves of elevation through time for each of the selected clades, whereby we included the elevational history of the Andean ranges that fall within the distribution ranges of the clades (**Supplementary materials)**, resulting in curves for (1) the northern and north-central Andes (domains 1-3 of Fig. 1) for Aromobatidae, Centrolenidae, Dendrobatidae and Hemiphractidae, (2) the central and southern Andes (domains 4-7 of Fig. 1) for Liolaemidea, and (3) the whole Andes (all 7 domains) for Leptodactylidae (**Fig. 2**).

**Figure 2.**
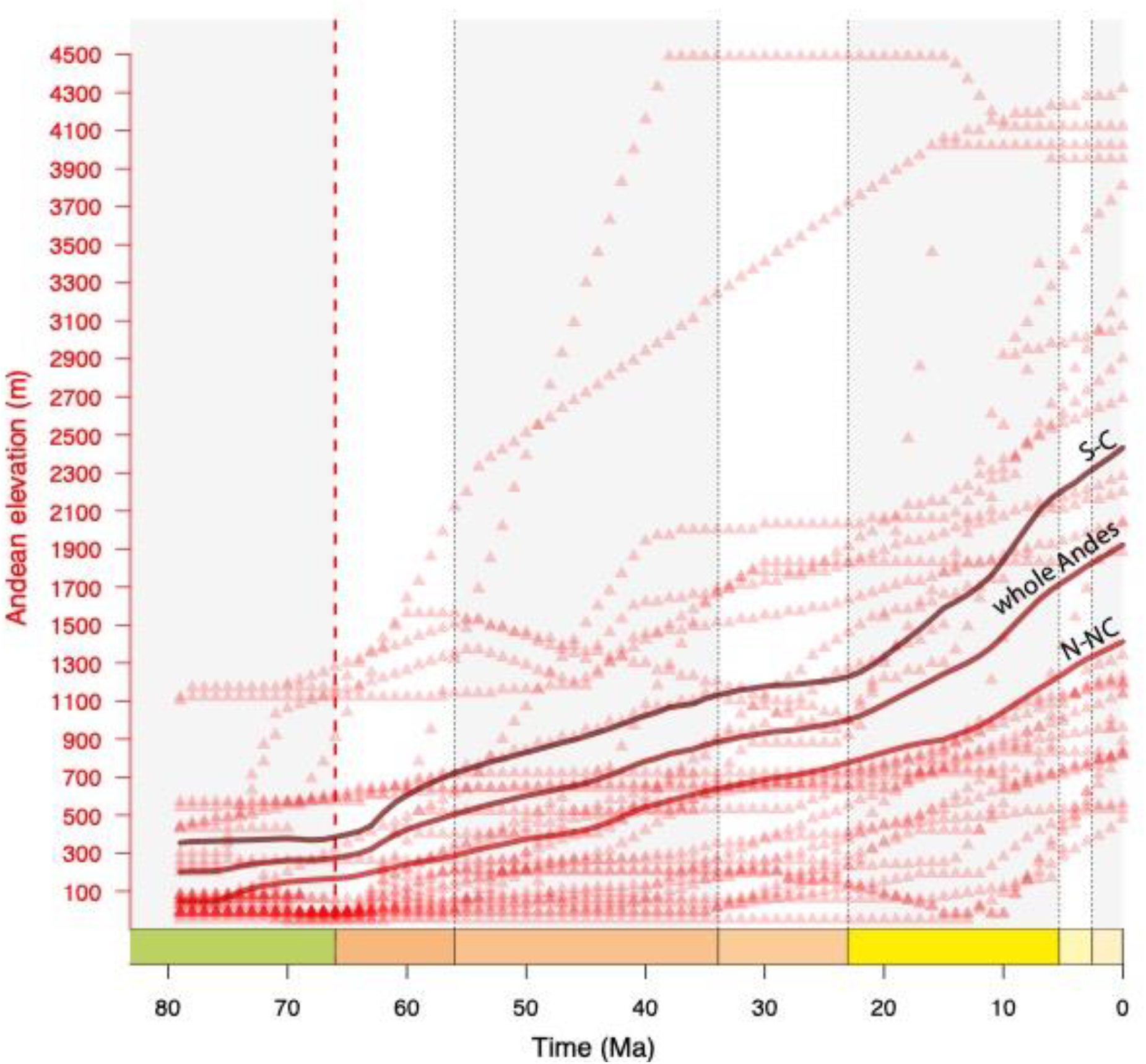
The Andean elevation curves used to estimate Neotropical diversification over the last 80 Myr. Red triangles represent data from individual geomorphological domains as reconstructed by Boschman (2021); solid lines represent the smoothed curves that are used as input in the elevation-dependent birth-death models (see descriptions in 2.4 and 2.5). S-C: southern and central Andes (domains 4-7 of Fig. 1), paleoelevation curve for Liolaemidea; N-NC: northern and north-central Andes (domains 1-3 or Fig. 1), paleoelevation curve for Aromobatidae, Centrolenidae, Dendrobatidae and Hemiphractidae; whole Andes (all 7 domains of Fig. 1) curve for Leptodactylidae.

In the southern (Patagonian) Andes (**Fig. 1**, domain 7), uplift initiated the earliest, at around 100 Ma (Dalziel et al. 1974; Bruhn and Dalziel, 1977, Kohn et al. 1995, Fosdick et al. 2011; Horton 2018), and modern elevations were reached at around 55 Ma (Colwyn et al. 2019). During the Miocene, the southern part of the orogen experienced additional uplift, primarily through expanding its width (Davis et al. 1983; Blisniuk et al. 2005, 2006; Giambiali et al. 2016). In the central Andes, including the world’s second largest and highest plateau area (the Altiplano-Puna Plateau), significant uplift initiated along the western margin of the South American plate at around 70 Ma (Horton 2018). Since then, uplift has migrated eastward, with major phases of uplift in the late Paleocene-Eocene at the western margin of the plateau area (**Fig. 1**, domain 4), in the Miocene at the eastern margin of the plateau area (domain 5), and active uplift since the middle Miocene in the easternmost Subandean zone (domain 6)(Carrapa et al. 2006, 2014; Scheuber et al. 2006; Uba et al. 2006; Leier et al. 2013; Canavan et al. 2014; Garzione et al. 2014; Fiorella et al. 2015; Quade et al, 2015; Rohrman et al. 2016). Mountain building in the northern Andes resulted from ∼70 Ma collision of the leading edge of the Caribbean Plate with the South American continental margin at the latitude of what is today Ecuador, and occurred initially only at the location of collision (Montes et al. 2019).Most of the northwestern corner of the South American continent remained below sea level throughout the Late Cretaceous and early Paleogene (Sarmiento-Rojas 2019). During the Paleogene, uplift slowly migrated to the north and east, reaching the Central Cordillera (part of domain 1) in the Eocene and the Perija Range and Santander Massif in the latest Eocene-Oligocene (part of domain 2). During the early Miocene, the Eastern Cordillera, Garzon Massif (part of domain 2) and Merida Andes (domain 3) experienced pronounced uplift, and uplift in these latter eastern ranges intensified since the Miocene (Gómez et al. 2005a,b; Villagómez et al. 2011; Anderson et al. 2015, 2016; Bermúdez et al. 2017; Horton 2018).

### 2.3. Temperature data

To capture the major trends in global climate through time throughout the clades’ evolutionary histories, we computed a temperature curve based on δ18O data measured from deep-sea benthic foraminifera shells preserved in oceanic sediments (**Fig. 3**). For the Cenozoic, we used the dataset from Westerhold et al. (2020; available at: https://doi.org/10.1594/PANGAEA.917717), and for the Cretaceous (80-66 Ma), the dataset from Veizer and Prokoph (2015; available at: https://doi.org/10.1016/j.earscirev.2015.03.008). To convert δ18O measurements to temperature values, we used the equations of Hansen et al. (2013), which convert δ18O to deep-ocean temperatures (Tdo) and subsequently, to surface temperatures (Ts); these equations are provided in the supplementary materials 1. We then summarized these data into a continuous estimate of temperature through time through calculation of a smoothing spline (degrees of freedom: 80). While each individual data point is subject to certain biases (e.g. some of them do not account for sea-level fluctuations, which are important during periods of large-scale glaciations, Cramer et al. 2011), the spline curve smoothens such biases, as well as geographical variations, providing a reliable estimate of global temperature trends (Veizer and Prokoph 2015). The surface temperature curve reflects planetary-scale climatic trends that can be expected to have led to temporally coordinated diversification changes in several clades rather than local or seasonal fluctuations (Erwin 2009; Hannisdal and Peters 2011; Mayhew et al. 2012; Condamine et al. 2019a).

**Figure 3.**
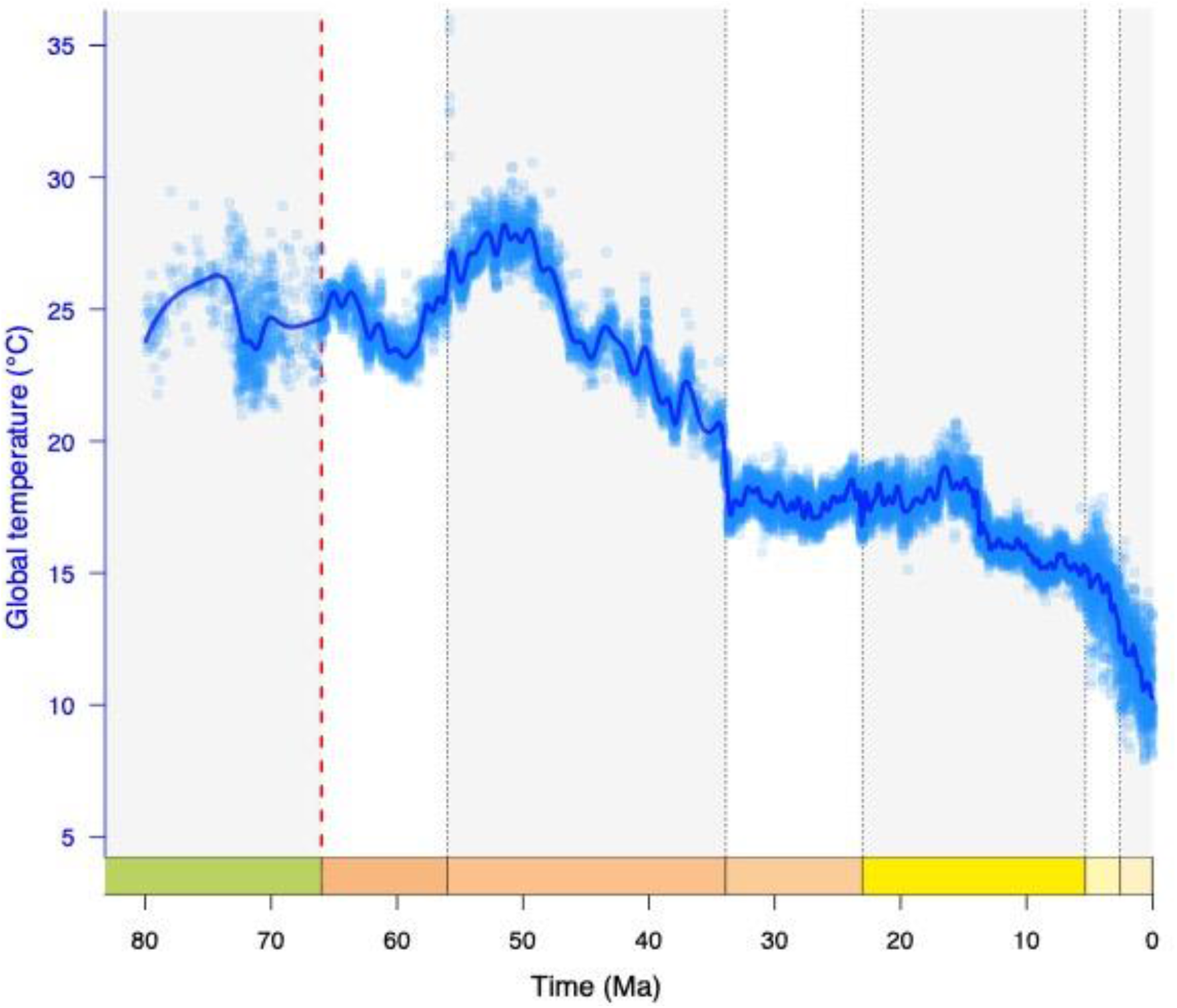
The global temperature curve used to estimate Neotropical diversification over the last 80 Myr. The trend of temperature is based on oxygen isotope ratios in benthic foraminifera shells; data from Veizer and Prokoph (2015) for the Late Cretaceous, and Westerhold et al. (2020) for the Cenozoic. Isotope ratios are converted to surface temperatures using the equations from Hansen et al. (2013). Blue circles represent data from individual δ^18^O measurements; the solid line represents the smoothed curve that is used as input in the temperature-dependent birth-death models (see descriptions in 2.4 and 2.5).

### 2.4. Modelling the effects of environmental change on diversification

Many studies have shown that the environment, shaped by long-term climate change and plate tectonics, plays a prominent role in the diversification of species over evolutionary and geological time scales (e.g. Mayhew et al. 2012; Zaffos et al. 2017). Yet, these studies have mostly based their conclusions on descriptive comparisons between dated speciation or extinction events estimated from fossil or phylogenetic data and paleoenvironmental curves (e.g. Delsuc et al. 2004; Antonelli et al. 2009; Eronen et al. 2015; Fan et al. 2020). Such an approach is suitable for events that occurred “instantaneously” in the context of geological time (i.e. within a million year), such as mass extinctions, but may not always be appropriate when comparing slower environmental changes. Most climate “events”, such as the Eocene Climatic Optimum (∼56-46 Ma), the middle Miocene climatic optimum (∼17-15 Ma) or the overall Cenozoic cooling trend, take place over longer timescales (Veizer and Prokoph 2015; Westerhold et al. 2020), and so do environmental changes related to plate tectonics such as the uplift of the Andes, which took place over the last ∼100 Myr (Boschman 2021). Furthermore, geological events are often intrinsically related to climate change, which complicates linking isolated environmental changes to single diversification events. For example, the uplift of the Andes affected climate in the eastern Pacific Ocean and above the South American continent (Sepulchre et al. 2010; Armijo et al. 2015), and uplift of the Qinghai-Tibetan Plateau shaped the Asian monsoon system (Molnar et al. 1993; An et al. 2011; Favre et al. 2015). In addition, the aforementioned descriptive approaches generally use phylogenetic trees — branching trees that represent the evolutionary relationships among species — that have uncertainties in their configuration and in age estimates of the speciation events, which makes comparison between a given environmental variable and the diversification processes ambiguous. Finally, these approaches do not quantify the long-term effect of environmental factors in shaping diversification rates, but rather look for punctuated change in diversification.

To account for these issues, methods have been developed in recent years to quantitatively explore factors that are potentially linked to speciation and/or extinction of lineages throughout geological and evolutionary time, thereby providing new opportunities to address questions about the mechanisms that shape diversity patterns (Egan et al. 2008; Condamine et al. 2013; Davis et al. 2016). Such methods have been made possible by the increased availability of statistical tools to analyse molecular and fossil data, and accessibility of time-calibrated molecular phylogenies. We focus in this study on phylogeny-based approaches, but note that these are beginning to converge with fossil-based approaches in their conceptual development and inferences to estimate temporal variations of diversification rates (Morlon et al. 2011; Silvestro et al. 2014, 2019), and evolutionary response to variations of the environment (Condamine et al. 2013; Silvestro et al. 2015; Lehtonen et al. 2017).

The approach used in this study, developed by Condamine et al. (2013, 2019a) and hereafter termed the environment-dependent diversification model, builds on time-dependent diversification models (Nee et al. 1994; Morlon et al. 2011), but allows speciation and extinction rates to depend not only on time but also on an external variable (which may vary through time). This methodology is implemented in the R-package *RPANDA* (Morlon et al. 2016). Clades are assumed to evolve under a birth-death process, in which both speciation and extinction follow a Poisson process, meaning that the expected waiting time to an event follows an exponential distribution (Nee 2006). As a result, we assume the speciation and extinction functions to be exponential. Speciation (λ) and extinction (μ) rates can vary through time, and both can be influenced by one or several environmental variables (here: temperature (*T)*, and Andean elevation *(A)*) that also vary through time. We consider the phylogeny of *n* species sampled from the present, and allow for the possibility that some extant species are not included in the sample by assuming that each extant species was sampled with probability *f* ≤ 1. Time is measured from the present to the past such that it denotes branching times in the phylogeny.

### 2.5. Analyzing the diversification of Andean clades

In this study, we fitted 14 diversification models to each of the six selected phylogenies using maximum likelihood (Stadler 2013; Morlon 2014). We consider four types of models with diversification rates that are constant (2 models), time-varying (4 models), temperature-dependent (4 models), and elevation-dependent (4 models) (**Fig. 4A; Table 2**). These models are fitted by maximum likelihood using the *fit_bd* (for the time-constant and time-varying models) and *fit_env* functions (for the temperature- and elevation-dependent models) from the R-package RPANDA 1.9 (Morlon *et al*. 2016). We accounted for missing species by specifying the sampling fraction corresponding to each phylogeny. We used the “crown” condition, which conditions the likelihood of a speciation event at the crown age and survival of the two daughter lineages.

**Table 2.** Descriptions of the 14 different birth-death models fitted to the six Neotropical amphibian and squamate families, including speciation and extinction equations, the number of free parameters to optimize by maximum likelihood, and the acronym as used in Table 3, which reports the results per family.

| Type of model | Model description | Model equation | Number of parameters | Model acronym in Table 3 |
| --- | --- | --- | --- | --- |
| Constant-rate models | Constant speciation and no extinction | $\lambda(t)=\lambda_0$ and $\mu(t)=0$ | 1 | BCST |
| | Constant speciation and constant extinction | $\lambda(t)=\lambda_0$ and $\mu(t)=\mu_0$ | 2 | BCSTD CST |
| Time-dependent models | Speciation variable and no extinction | $\lambda(t)=\lambda_0 e^{\alpha t}$ and $\mu(t)=0$ | 2 | BTimeVar |
| | Speciation variable and constant extinction | $\lambda(t)=\lambda_0 e^{\alpha t}$ and $\mu(t)=\mu_0$ | 3 | BTimeVarDCST |
| | Constant speciation and extinction variable | $\lambda(t)=\lambda_0$ and $\mu(t)=\mu_0 e^{\beta t}$ | 3 | BCSTDTimeVar |
| | Both speciation and extinction variable | $\lambda(t)=\lambda_0 e^{\alpha t}$ and $\mu(t)=\mu_0 e^{\beta t}$ | 4 | BTimeVarDTimeVar |
| Temperature-dependent models | Speciation variable and no extinction | $\lambda(t)=\lambda_0 e^{\alpha T(t)}$ and $\mu(t)=0$ | 2 | BTempVar |
| | Speciation variable and constant extinction | $\lambda(t)=\lambda_0 e^{\alpha T(t)}$ and $\mu(t)=\mu_0$ | 3 | BTempVarDCST |
| | Constant speciation and extinction variable | $\lambda(t)=\lambda_0$ and $\mu(t)=\mu_0 e^{\beta T(t)}$ | 3 | BCSTDTempVar |
| | Both speciation and extinction variable | $\lambda(t)=\lambda_0 e^{\alpha T(t)}$ and $\mu(t)=\mu_0 e^{\beta T(t)}$ | 4 | BTempVarDTempVar |
| Elevation-dependent models | Speciation variable and no extinction | $\lambda(t)=\lambda_0 e^{\alpha A(t)}$ and $\mu(t)=0$ | 2 | BAndesVar |
| | Speciation variable and constant extinction | $\lambda(t)=\lambda_0 e^{\alpha A(t)}$ and $\mu(t)=\mu_0$ | 3 | BAndesVarDCST |
| | Constant speciation and extinction variable | $\lambda(t)=\lambda_0$ and $\mu(t)=\mu_0 e^{\beta A(t)}$ | 3 | BCSTDAndesVar |
| | Both speciation and extinction variable | $\lambda(t)=\lambda_0 e^{\alpha A(t)}$ and $\mu(t)=\mu_0 e^{\beta A(t)}$ | 4 | BAndesVarDAndesVar |

**Figure 4.**
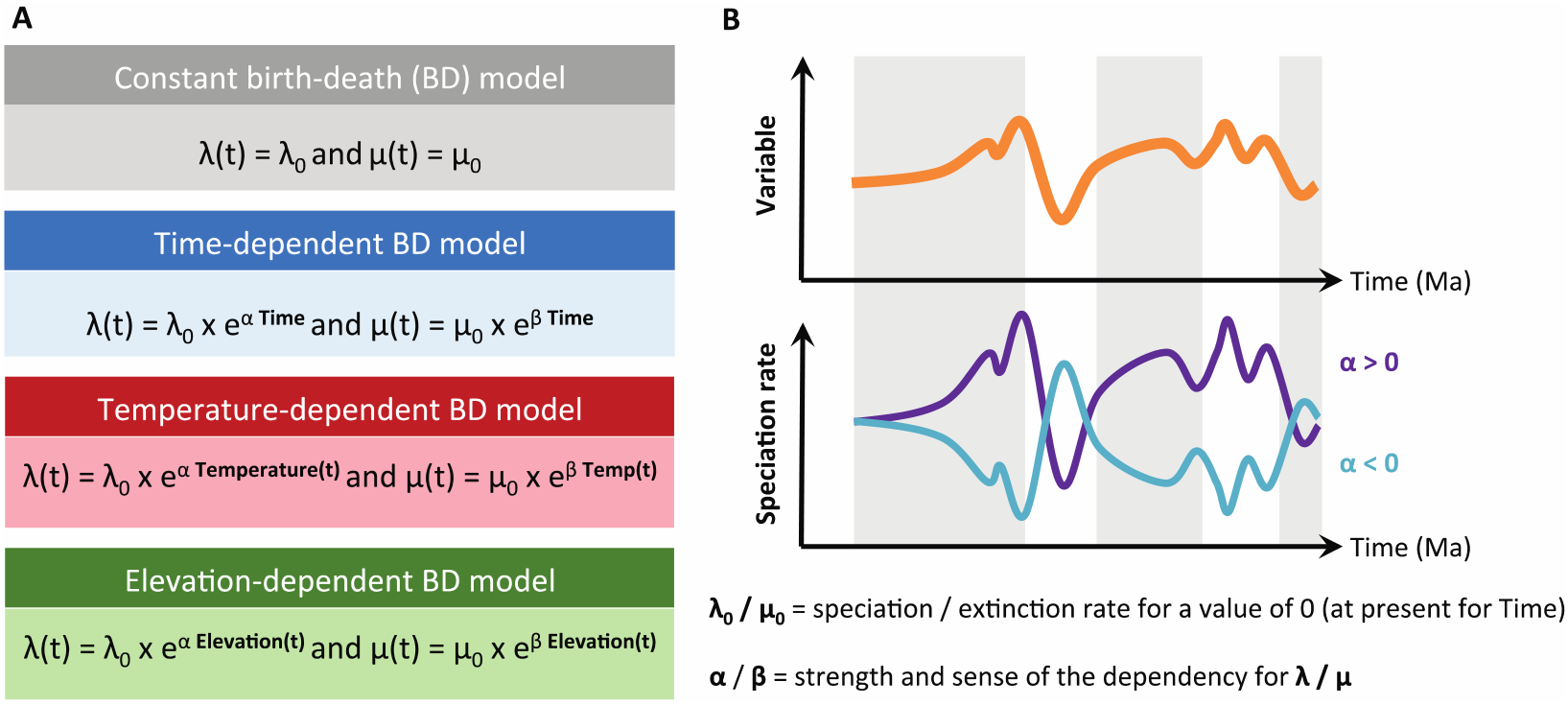
Schematic description of the birth-death (BD) models used to estimate diversification rates. A) List of the four types of BD models with corresponding rate functions for speciation and extinction. B) Example of speciation rate through time λ(*t*) obtained from the relationship between speciation rate and the past variation of an environmental variable (orange curve). The relationship between speciation rate and environment can be positive (α > 0; purple curve) or negative (α < 0; light blue curve). If α = 0 there is no effect of the environment, and the speciation rate is constant.

In the time-dependent models, λ or both λ and μ vary as a continuous function of time (**Table 2**): λ(t)=λ0e^αt^ or μ(t)=μ0e^βt^, where λ0 (μ0) is the speciation (extinction) rate at present. A positive α (β) reflects a slowdown of speciation (extinction) towards the present, while a negative α (β) reflects a speed-up of speciation (extinction) towards the present, and the sign and value of α and β depends on the data and model optimization. In the environment-dependent models, speciation rates, extinction rates, or both vary as a continuous function of Andean elevation *A* (**Fig. 2**) or temperature *T* (**Fig. 3**), for which the curves are computed using a spline interpolation which the degree of freedom set to 80 (df=80 in the *fit_env* function). We consider the same exponential dependency as above, but with *t* replaced by *T(t)* or *A(t)*. In this case λ0 (μ0) is the expected speciation (extinction) rate under a temperature of 0°C or an altitude of 0 meter, and α (β) measures the sign and strength of the temperature or paleoelevation dependence (**Fig. 4B**). A positive α (β) indicates that speciation (extinction) rates are higher under warm climatic conditions or when elevations were high, while a negative α (β) indicates that speciation (extinction) rates are higher under cold climatic conditions or when elevations were low (**Fig. 4B**).

We fitted each of the models to each phylogeny by maximum likelihood, starting with the simplest (constant rate) models and progressively increasing in complexity. The maximum-likelihood algorithm optimizes parameter values (of λ0, μ0, α and/or β) that maximize the probability of the observed data (the phylogenetic tree) under a given model. Because these optimization algorithms can be sensitive to the choice of initial parameter values (they can converge to local optima in the vicinity of the initial parameter values), we informed the initial parameter values of more complex models by those previously estimated on simpler models. The 14 tested models are not all nested, and we used the corrected Akaike Information Criterion (AICc; Burnham and Anderson 2002) to compare models. The AICc is useful to compare the probability of observing branching times as explained by various individual paleoenvironmental variables (Condamine et al. 2015, 2018, 2019a). A series of models can be designed to quantify the effect that various environmental variables, taken in isolation, had on diversification. We thus compared the AICc scores for the best-fit models between multiple environmental variables (temperature, Andean uplift) to determine which has the strongest effect on diversification. The best-fitting model is selected using Akaike weights (AICω).

## 3. Results

Within the six clades, we found that three clades primarily supported an environment-independent model of diversification (**Figs. 5 and 6, Table 3**): the constant-rate model best fitted the diversification of Dendrobatidae (AICω=0.523; AICω=0.286 for the second best model that is a temperature-dependent model, **Fig. 5E, F**) and of Hemiphractidae (AICω=0.447; AICω=0.288 for the second best model that is an elevation-dependent model, **Fig. 6A, B**), while the time-dependent model best explains the diversification of Liolaemidae (AICω=0.447; AICω=0.394 for the second best model that is an elevation-dependent model, **Fig. 6E, F**). The remaining three clades showed statistical support for an environment-dependent model of diversification with either a temperature-dependent model for Centrolenidae (AICω=0.882; AICω=0.096 for the second best model that is an elevation-dependent model, **Fig. 5C, D**), or an elevation-dependent model for Aromobatidae (AICω=0.549; AICω=0.286 for the second best model that is a time-dependent model, **Fig. 5A, B**), and Leptodactylidae (AICω=0.536; AICω=0.279 for the second best model that is a constant-rate model, **Fig. 6C, D**).

**Table 3.**
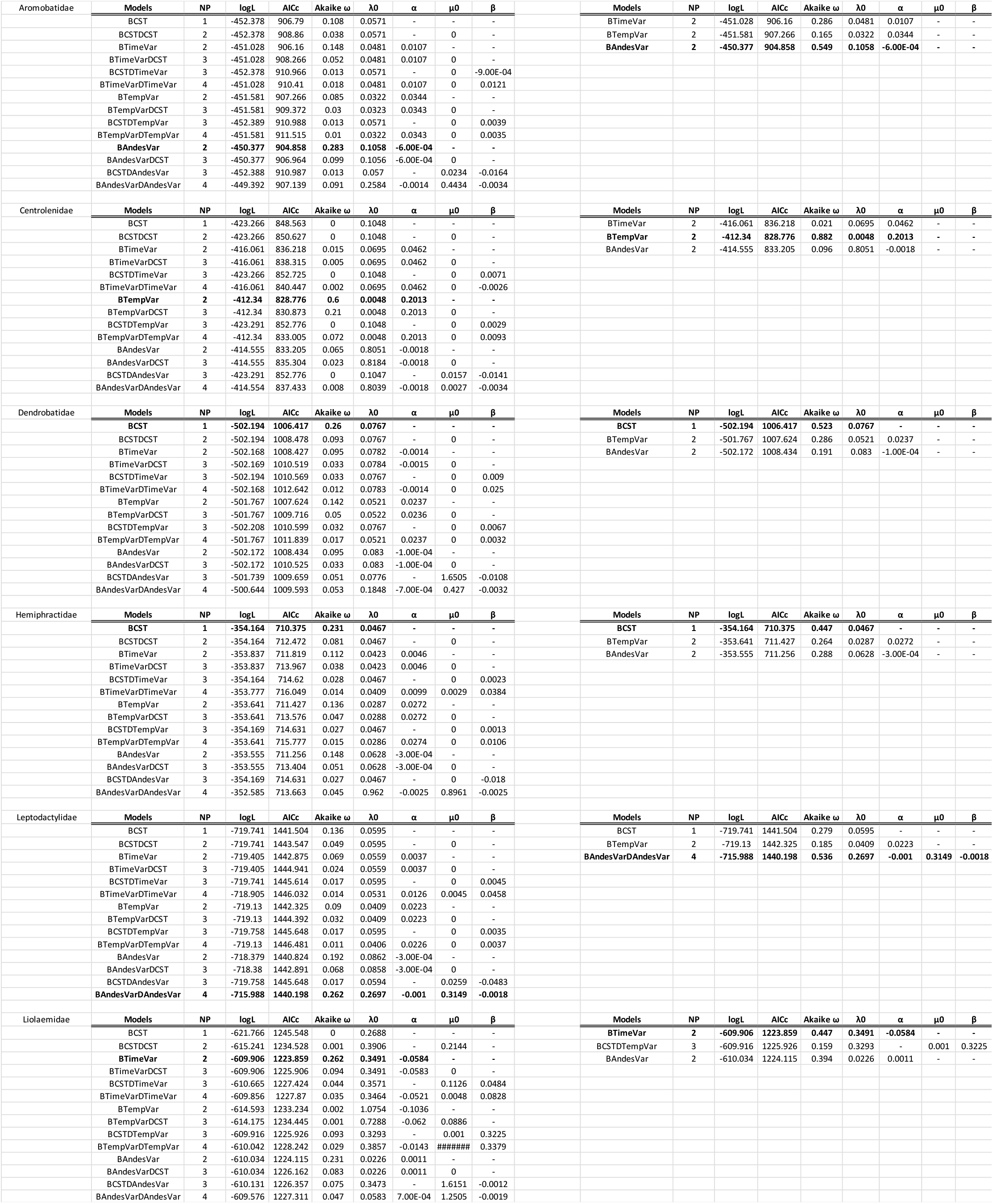
Results of all diversification analyses performed on the six Neotropical amphibian and squamate families. Panels on the left report the models (14 in total), the number of parameters in each model, the estimated log-likelihood (logL), the corrected Akaike information criterion (AICc), the Akaike weight of the model considering all 14 models (AICω), and the corresponding parameter estimates (λ0 = speciation rate at present, α = parameter controlling the dependency of speciation rate on time or temperature, μ0 = extinction rate at present, and β = parameter controlling the dependency of extinction rate on time or temperature). The best-fitting model is highlighted in bold. Panels on the right report the same, except the Akaike weight (AICω) is now calculated after considering only the best-fit models of each of the three categories (environment-independent, temperature-dependent, elevation-dependent).

**Figure 5.**
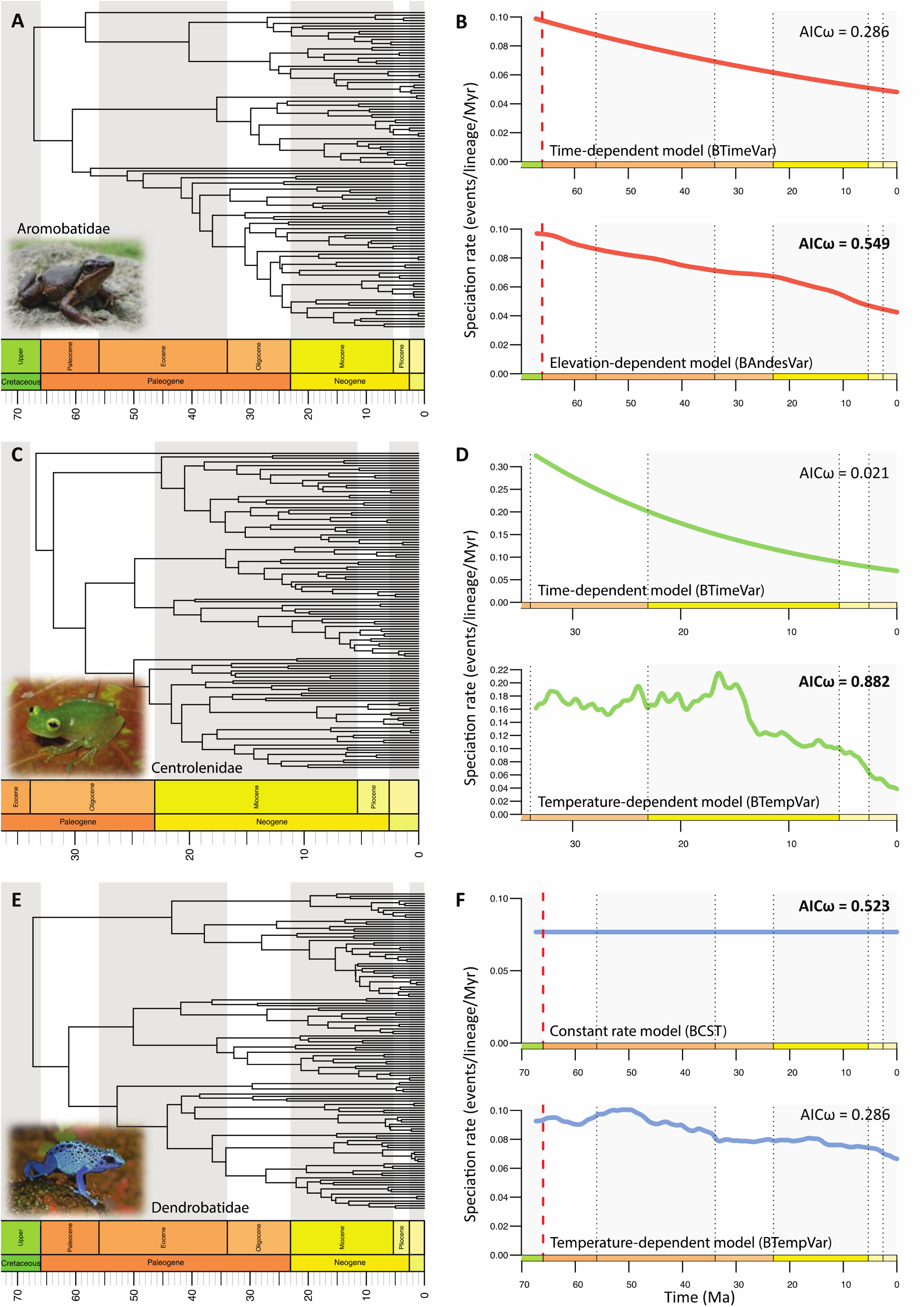
Estimating the diversification processes for three Neotropical frog families distributed in the Andes. Left panels show the time-calibrated phylogenies for three Neotropical frog families: A) Aromobatidae (cryptic forest frogs), C) Centrolenidae (glass frogs), and E) Dendrobatidae (poison dart frogs); from Hutter et al. (2017). Right panels show, for each family, the results of the best environment-independent diversification model (top panel) and of the best environment-dependent diversification model (bottom panel). Akaike weights (AICω) for both models are shown and the best-fit model for each family is indicated in bold. Pictured for Aromobatidae is *Rheobates palmatus* by G.A. Chaves Portilla, for Centrolenidae is *Hyalinobatrachium fleischmanni* by M. Rivera Correa, and for Dendrobatidae is *Dendrobates tinctorius* by H. Zell.

**Figure 6.**
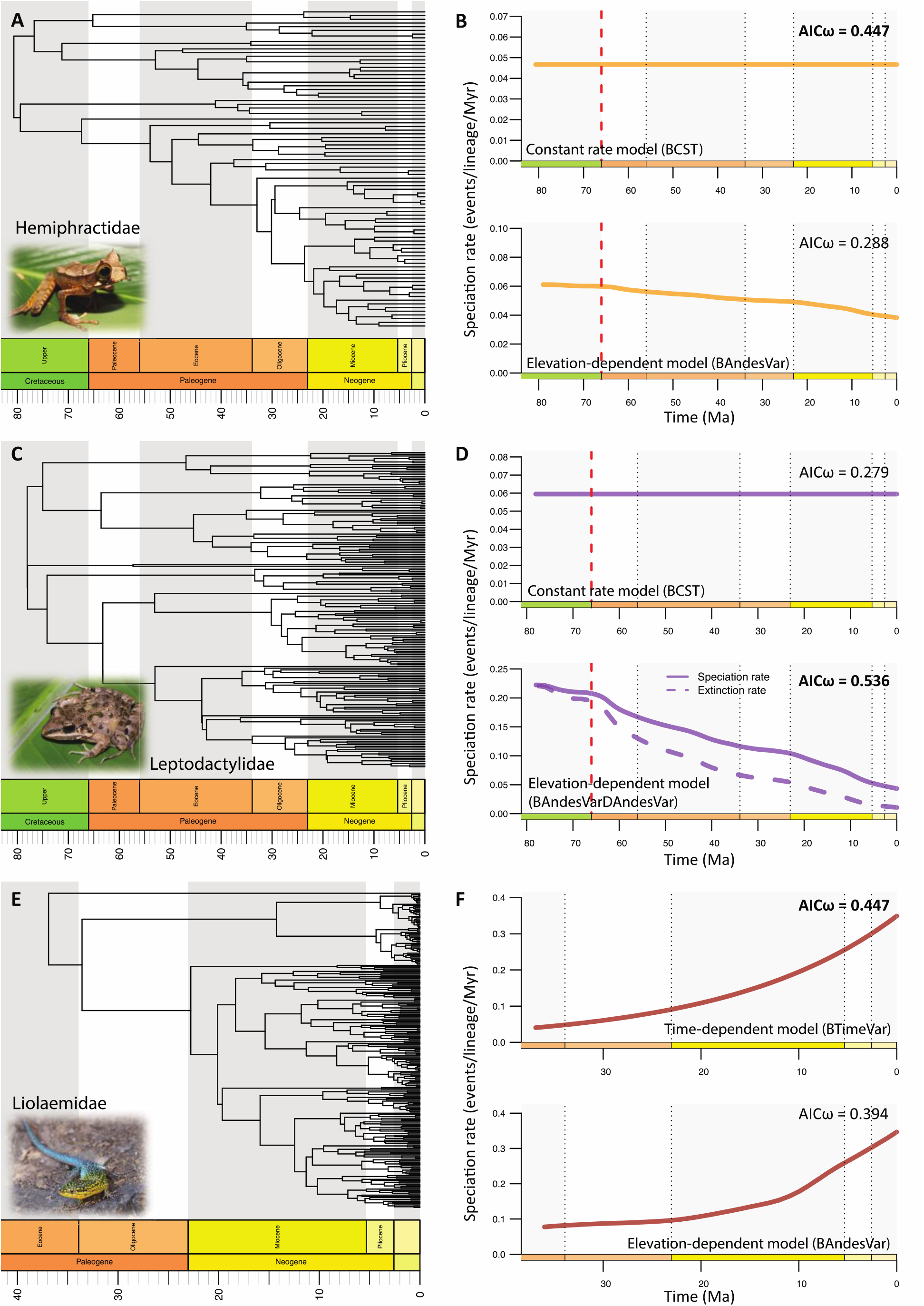
Estimating the diversification processes for two Neotropical frog families and one Neotropical lizard family distributed in the Andes. Left panels show the time-calibrated phylogenies for three Neotropical frog families: A) Hemiphractidae (marsupial frogs), C) Leptodactylidae (southern frogs), and E) Liolaemidae (liolaemid lizards); from Hutter et al. (2017) and Esquerré et al. (2019). Right panels show, for each family, the results of the best environment-independent diversification model (top panel) and of the best environment-dependent diversification model (bottom panel). Leptodactylidae is the only clade for which extinction rates are estimated in the best-fit model, which are shown in panel D. Akaike weights (AICω) for both models are shown and the best-fit model for each family is indicated in bold. Pictured for Hemiphractidae is *Gastrotheca cornuta* by B. Gratwicke, for Leptodactylidae is *Leptodactylus fragilis* by E. Alzate, and for Liolaemidae is *Liolaemus tenuis* by Kaldari.

For Centrolenidae, the temperature-dependent model estimated a positive correlation between speciation rates and global temperatures (α=0.2013, **Fig. 5D**), which translates into faster speciation during warmer periods. For the other five clades, the temperature-dependent model was not selected as the best-fit model, but did indicate a positive correlation between speciation rates and global temperatures, except for Liolaemidae (**Table 3**). The elevation-dependent model estimated a negative correlation between speciation rates and Andean elevation for both Aromobatidae and Leptodactylidae (α=-6.00E-04, -0.001, respectively, **Figs. 5B and 6D**). This implies that these groups diversified faster in the early stages of the Andean orogeny. Moreover, the model for Leptodactylidae indicated a negative correlation between extinction rates and Andean elevation (β=-0.0018), which suggests higher extinction rates during the initial phase of mountain building.

Except for Liolaemidae, all elevation-dependent models (whether best-fit or not) estimated negative correlations between speciation rates and Andean elevation (**Table 3**), meaning high speciation rates during low elevations, and thus, during the early stages of the Andean orogeny. For Liolaemidae, one of the two younger (∼37 Ma) clades, the best fit-model is a time-dependent model, which estimated an increase in speciation through time (α=-0.0584) with a speciation rate of λ0=0.3491 events/lineage/Myr at present (**Fig. 6F**). However, it is important to mention the second best-fitting model indicating a positive correlation (α=0.0011) between speciation and Andean elevation, because the difference between the two best models is small (ΔAICc=0.256, **Table 3**). Liolaemidae is the only studied clade showing a positive relationship between speciation and Andean elevation (**Fig. 6F**). This result suggests higher speciation rates during the later stages of Andean orogeny, when the Andes were higher.

The support for the models best explaining the diversification of the six clades is variable (AICω ranging from 0.231 to 0.60, with a mean of 0.316; **Table 3**), but always above the 0.071 weight (1/14) that would be expected if all models were equally likely. Nonetheless, after selecting the best-fitting model from each of the three main types (non-environment-dependent [including constant and time-dependent], temperature-dependent or elevation-dependent), and comparing only these three, the strength of support of the best-fitting models was reasonably high (mean AICω of 0.473 for non-environment-dependent models, 0.882 for temperature-dependent models, and 0.542 for elevation-dependent models). These values are consistently above the 0.33 (1/3) weight that would be expected if non-environment-dependent, temperature-dependent rate and elevation-dependent models models were equally likely.

## 4. Discussion

### 4.1 Long-lived mountain building triggering both ancient and recent radiations

We have studied potential drivers behind the diversification histories of six South American clades, four originating in the Late Cretaceous, and two during the latest Eocene-earliest Oligocene. We show that for Dendrobatidae, Hemiphractidae and Liolaemidae, the diversification history is likely primarily driven by environment-independent factors; for Centrolenidae, primarily by Cenozoic temperature variations, and for Aromobatidae and Leptodactylidae, primarily by Andean mountain building (**Figs. 4 and 5**). Interestingly, the two clades for which we link diversification to Andean elevation (Aromobatidae and Leptodactylidae) diversified faster during the incipient (Late Cretaceous - Paleogene) phase of Andean orogeny. During this phase, the western margin of the South American continent transformed from a continental margin hosting a magmatic arc with localized topography only and average elevations below 1,000 m (Vergara et al. 1995; Boekhout et al. 2012; Horton 2018), into a fold-thrust belt with significant and continuous topography (Horton 2018; Boschman 2021; **Fig. 1**). During this time, the Andes were still significantly lower and less wide compared to their modern extent, but the first (arid) high-altitude habitats were appearing, especially in the southern (Mathiasen and Premoli 2010) and central (Quade et al. 2015) Andes.

Aromobatidae and Leptodactylidae are both widely distributed in the Andes, but for both clades, the center of species richness is located outside of the Andes. Nonetheless, for Aromobatidae, diversification has previously been linked to early Miocene uplift of the Eastern Cordillera of Colombia and the resulting middle Miocene onset of aridity in the Magdalena Valley (Muñoz-Ortiz et al. 2015). Additionally, indirect effects of Andean uplift shaped river and lake system dynamics in western Amazonia, which acted as a source of aromobatid diversity for the rest of Amazonia (Vacher et al. 2017; Réjaud et al. 2020). With this study, we now add evidence for the influence of early Andean orogeny to Aromobatidae and Leptodactylidae diversification (**Figs. 4B and 5D**). This result is in line with results from Santos et al. (2009), who concluded that montane habitats in the Guiana Shield and Atlantic Forest regions were colonized at a later stage.

Although diversification of Liolaemidae is best explained by an increase of speciation toward the present without invoking the role of environment, the second best-fit model (almost equally strong; **Fig. 5F**) indicates a positive relation between speciation rates and elevation. This result is in line with the study of Esquerré et al. (2019), who concluded that uplift of the Andes promoted Liolaemidae diversification through allopatric fragmentation, and acted as a species pump for the surrounding regions.

Perhaps surprisingly, our analyses did not select the elevation-dependent model as the best fit model for the hyloid frog clades Centrolenidae, Dendrobatidae and Hemiphractidae, contrary to previous conclusions reached concerning these groups (Hutter et al. 2017). For Centrolenidae, our analyses instead yielded strong support for the temperature-dependent model with high speciation rates during warm periods (**Fig. 4D, F**). This suggests a substantial effect of Cenozoic temperature variations in shaping Neotropical diversification in the Andes; a model that had so far not been tested. Alternatively, these results could be misleading, because the temperature-model has been shown to fit well overall across tetrapod families (Condamine et al. 2019a). Finally, the best model for Dendrobatidae and Hemiphractidae is a model with constant speciation rates through time. For the latter clade, this model could reflect true evolutionary mechanisms, because marsupial frogs are unique among anurans in that the eggs develop on the back or in dorsal pouches in the female (Castroviejo-Fisher et al. 2015). Females of *Cryptobatrachus, Hemiphractus*, and *Stefania* carry their eggs and young on their backs, and in other hemiphractids, the eggs develop in a closed pouch (*Gastrotheca* and some *Flectonotus*) or in an open pouch (other *Flectonotus*). This breeding behaviour could buffer rates of diversification because these species are independent of water sources; they avoid the eggs having to hatch in ponds and develop as free-swimming tadpoles, which is prone to predation, and relies on favorable environmental conditions.

The effect of Paleogene orogeny that we report differs from the conclusions of a wealth of studies finding recent and rapid mountain clade radiations across the world’s mountain belts (e.g. Lenevue et al. 2009; Mao et al. 2010; Drummond et al. 2012; Jabbour and Renner 2012; Wang et al. 2012; Schwery et al. 2015; Lagomarsino et al. 2016; Esquerré et al. 2019; Ye et al. 2019). These findings do not contradict ours, but rather, it shows that so far, ancient mountain radiations have largely been undetected. In the study of Andean radiations, two of the main reasons for the absence of ancient radiations are the focus on Neogene Neotropical clades, and in some cases, the misconception that the bulk of Andean topography is young. By incorporating the early history of mountain building, and selecting old (Late Cretaceous) clades, we demonstrate the potential effect of the initial emergence of high-altitude habitats on diversification. This implies that throughout the long-lived history of surface uplift in the Andes, mountain building drove the diversification of different clades at different times, while not affecting yet other clades altogether.

An analogy can be made with the history of uplift and diversification in the Tibetan Plateau region. Renner (2016) showed that many macroevolutionary and biogeographic studies have linked (potentially non-existent) recent uplift to young and rapid radiations (e.g. Favre et al. 2015, 2016; Ebersbach et al. 2017; Ye et al. 2019), despite geological evidence pointing to an Eocene 4 km-high Tibetan Plateau. Although controversy about the uplift history of the Tibetan Plateau is not yet fully resolved (see e.g. Rowley and Garzione 2007; Botsyun et al. 2019), both the synthesis of Renner (2016) and the current study illustrate the importance of accurate and detailed paleogeographic reconstructions that capture the complexity and heterogeneity of mountain building over geological time, and the importance of connecting geologists and biologists in the attempt to solve these interdisciplinary problems.

### 4.2 Limitations of current modelling approaches and future perspectives

Distinguishing between the drivers of diversification requires datasets that adequately sample large geographical and taxonomic scales through both long and short time intervals (e.g. Marx and Uhen 2010; Eronen et al. 2015; Lagomarsino et al. 2016; Lehtonen et al. 2017; Condamine et al. 2018, 2019). To unravel the processes governing the tempo and mode of lineage diversification, phylogenies are required that encompass at least 80% of the total standing diversity within the clade of interest (e.g. Cusimano and Renner 2010; Höhna et al. 2011; Davis et al. 2013). This sampling ideally captures the entire range of ecological, geographic and morphological diversity, which requires accurate taxonomy, and clear documentation of the numbers of identifiable species in the region(s) of interest. As taxonomists remain underfunded, consensus on this basic information is not yet reached for all groups, yet is increasingly available for vertebrate groups. Knowledge of species ecology, including species interactions and species traits, may greatly enhance the design of macroevolutionary analyses in relation with hypotheses to test. Fortunately, phylogenies are increasingly improving, even in species-rich groups, and biological data, including morphology, ecology and distribution, are accumulating at an unprecedented rate (e.g. Pigot et al. 2020).

The state-of-the-art analytical framework used in this study is a step forward in our understanding of the diversification of lineages and its drivers, but is not without limitations. First, as with many macroevolutionary models, it suffers from the fundamental dilemma of extrapolation of correlations to causations. Second, difficulties remain in the estimation of speciation and extinction rates from phylogenies of extant species alone (e.g. Nee 2006; Ricklefs 2007; Rabosky and Lovette 2008; Crisp and Cook 2009; Quental and Marshall 2010; Burin et al. 2019; Pannetier et al. 2021). For instance, simulation and empirical studies have shown that it is complicated to distinguish between decreasing speciation or increasing speciation when the net diversification rate declines through time (Rabosky and Lovette 2008; Crisp and Cook 2009). Furthermore, Burin et al. (2019) show that two popular methods perform equally well when varying speciation rates control decline, but when decline was only caused by an increase in extinction rates both methods wrongly assign the variation in net diversification to a drop in speciation. Several clades of our study show decay of speciation through time and no extinction (**Figs. 5 and 6**), and we thus remain cautious with these estimations. Recently, Pannetier et al. (2021) demonstrated that phylogenetic trees contain insufficient information to detect the presence or absence of diversity-dependent diversification, and we thus refrained ourselves to use this model although it can provide additional tests of macroevolutionary hypotheses on the role of within-clade species interactions compared to abiotic factors (Condamine et al. 2019a). Third, it remains challenging to decipher the direct drivers of speciation or extinction rates and events (Ezard et al. 2016), as for instance, paleoelevation *per se* may not be a direct driver of diversification, but rather, the many indirect consequences of uplift, such as habitat fragmentation or local climate change (Lagomarsino et al. 2016; Pérez-Escobar et al. 2017; Nevado et al. 2018). Furthermore, we here compare environment-independent, temperature-dependent and elevation-dependent birth-death models, but temperature and elevation are not the only possible drivers of diversification, as many other a(biotic) processes (e.g. regional sea level variations, precipitation, erosion, interactions between species) may have played a role. However, compared to paleoelevation and global paleotemperature, quantifying or reconstructing these other possible drivers of diversification is a much bigger challenge, which hampers testing them in diversification models. In addition, it is questionable whether it truly is elevation (or temperature) that is driving diversification, rather than the change in elevation (or temperature). Although biological interpretation might become less straightforward, future studies might consider incorporating environmental-change diversification models, using the time-derivative of environmental variables.

To uncover the extent to which speciation and extinction vary and according to which drivers, macroevolutionary studies may need to combine diversification (birth-death) models with parametric biogeography models (Ree and Smith 2008; Quintero et al. 2015) or mechanistic eco-evolutionary simulation models (Rangel et al. 2018; Hagen et al. 2021). Such combined approaches can for example be used to estimate whether speciation occurred via allopatry (vicariance) or sympatry (ecological speciation), which specific factors in paleo-environmental change caused speciation and/or extinction, and in which locations these diversification events took place. It is worth mentioning that some models exist to test the role of diversity-dependence in the context of allopatry (Pigot et al. 2010; Valente et al. 2015).

New developments in the field of trait-dependent models form a second line of research contributing to our understanding of the interactions between biotic and abiotic factors and their effects on diversity dynamics (Maddison et al. 2007; Goldberg et al. 2011; Beaulieu and O’Meara 2016). Cantalapiedra et al. (2014) developed a climate-based trait-dependent diversification model, in which rates of speciation not only vary as a function of the same global temperature curve that we incorporated into our analysis, but also among traits. The model is thereby able to identify differences in diversification history between clades with different functional traits as a function of changes in environment. Cantalapiedra et al. (2014) showed that ruminant clades with three feeding modes (browsers, grazers and mixed feeders) diversified differently during warm or cold periods.

A remaining, outstanding question is which role competition plays in ecologically similar clades that are phylogenetically unrelated (Benton 1987; Hembry & Weber 2020). Although it is difficult to model the effect of external clades interacting with the focal clade and their effects on speciation and/or extinction with current models, some progress has been made with fossil-based models showing that clade competition occurred over long timescales (Liow et al. 2015; Silvestro et al. 2015). It is thus possible to assess the effect of competition on diversification, in which speciation and extinction rates are correlated with the diversity trajectory of another clade. Under competitive interaction scenarios, increasing species diversity has the effect of suppressing speciation rates and/or increasing extinction rates. Although in phylogenetic studies, diversity dependence is typically tested within a single clade (Etienne et al. 2012), our phylogeny-based diversification model can be extended to allow for competition among species that are not closely related, but share similar ecological niches (Condamine et al. 2020).

Lastly, studying the effects of environmental (i.e. climatic or geological) change requires accurate reconstructions of these changes through time. Despite the existence of contrasting datasets and resulting controversy, there is a relative abundance of data on the history of uplift of the two highest, largest (and most biodiverse) mountain belts on Earth: the Andes (see compilation in Boschman 2021) and the Tibetan Plateau-Himalayas-Hengduan mountains (e.g. An et al. 2001; Rowley and Garzione 2007; Botsyun et al. 2019), and to a lesser extent, of the North American Cordillera (e.g. Chamberlain et al. 2012). As a result, the majority of studies on environment-dependent diversification have focussed on these regions (Lagomarsino et al. 2016; Pérez-Escobar et al. 2017; Esquerré et al. 2019; Testo et al. 2019; Ding et al. 2020). Working towards a global understanding of the relationship between environmental change and lineage diversification (Antonelli et al. 2018a; Quintero and Jetz 2018; Rahbek et al. 2019a,b) calls for the collection of paleoelevation data on lesser-studied smaller orogens, particularly in species rich regions, such as for example New Guinea, hosting the world’s richest island flora (Cámara-Leret et al. 2020).

In conclusion, we still face important limitations in data availability as well as methodological shortcomings, but by acknowledging them we can target where to focus future efforts of the geological, macroevolutionary and biogeographical communities. Together, we may gain a better understanding of how environmental and biotic triggers are intertwined and of the rich, deep past of Earth’s biological diversity. We hope that our approach (and analytical framework) will help the forward movement in that direction, and that it will provide perspectives for future investigations on other model groups, and in other regions.

## 5. Conclusions

Analyzing the role of paleogeographic and paleoclimatic history in species diversification through macroevolutionary analyses has never been more exciting and promising than today. The conjunction of rapid and massive increases in the availability of biological datasets (phylogenies, fossils, georeferenced occurrences and ecological traits) and paleoclimate and paleogeographic reconstructions on the one hand, and the successful development of powerful analytical tools on the other, now enables assessing the relative roles of climate change and mountain building on lineage diversification.

Here, we analysed the diversification history of six Neotropical frog and lizard families in light of Cenozoic climate variations and Andean mountain building. The recent (Neogene) rise of the Andes is often considered as the prime driver of biological diversification in the Neotropics, but here we unveil a more complex evolutionary history for ancient yet species-rich frog and lizard families. We find clade-specific responses to temperature and elevation, and conclude that throughout the long-lived history of surface uplift in the Andes, mountain building drove the diversification of different clades at different times, while not directly affecting other clades. Although we find that the diversification of Aromobatidae and Leptodactylidae was influenced by the rise of Andes, we conclude that this effect played a role in the Late Cretaceous-Paleogene phase of orogeny, much earlier than previously proposed (i.e. Neogene). Moreover, we demonstrate that other drivers must be considered when studying mountain clade radiations: diversification rates of Centrolenidae were high during the warmer periods of the Cenozoic. Additionally, even a strict Andean-endemic lizard radiation shows mixed evidence for a direct role of Andean uplift driving the diversification. Therefore, our study argues that pre-Neogene environmental changes, either triggered by Andean uplift or resulting from global climate variations, should not be dismissed as drivers of Neotropical diversification, and calls for a greater appreciation of the ancient Andean paleo-elevation history in the build-up of species on the Earth’s most biodiverse continent.

## Data availability

All data files (phylogenies, environmental variables) and scripts necessary to perform the analyses will be provided via an online repository (Figshare).

## Acknowledgements

This work was funded by ETH postdoctoral fellowship 18-2 FEL-52, and by an “Investissements d’Avenir” grant managed by the Agence Nationale de la Recherche (CEBA, ref. ANR-10-LABX-25-01) and the ANR GAARAnti project (ANR-17-CE31-0009). We thank editor Daniele Silvestro, Pierre Sepulchre and an anonymous reviewer for very detailed and constructive comments, and Carina Hoorn for setting up this collaboration and the invitation to submit to this special issue.

## Supplementary materials

**Supplementary Table 1.** Equations (from Hansen et al., 2013, modified by Westerhold et al. 2020) to calculate deep-ocean temperature (T_do_) and surface air temperature (Ts) from δ^18^O data.

| $\delta^{18}O$ to $T_{do}$: | |
| --- | --- |
| 0.0 Ma to 3.660 Ma: | $T_{do} (^{\circ}C) = 1 - 4.4 * ((\delta^{18}O (‰) - 3.25 / 3)$ |
| 3.660 Ma to 34.025 Ma: | $T_{do} (^{\circ}C) = 5 - 8 * ((\delta^{18}O (‰) - 1.75 / 3)$ |
| 34.025 Ma to 67.0 Ma: | $T_{do} (^{\circ}C) = (-4 * \delta^{18}O (‰)) + 12$ |
| $T_{do}$ to $T_s$: | |
| 0.0 Ma to 1.81 Ma: | $T_s (^{\circ}C) = 2 * T_{do} + 12.25$ |
| 1.81 Ma to 5.33 Ma: | $T_s (^{\circ}C) = 2.5 * T_{do} + 12.15$ |
| 5.33 Ma to 67.0 Ma: | $T_s (^{\circ}C) = T_{do} + 14.15$ |

**Supplementary Figure 1.**
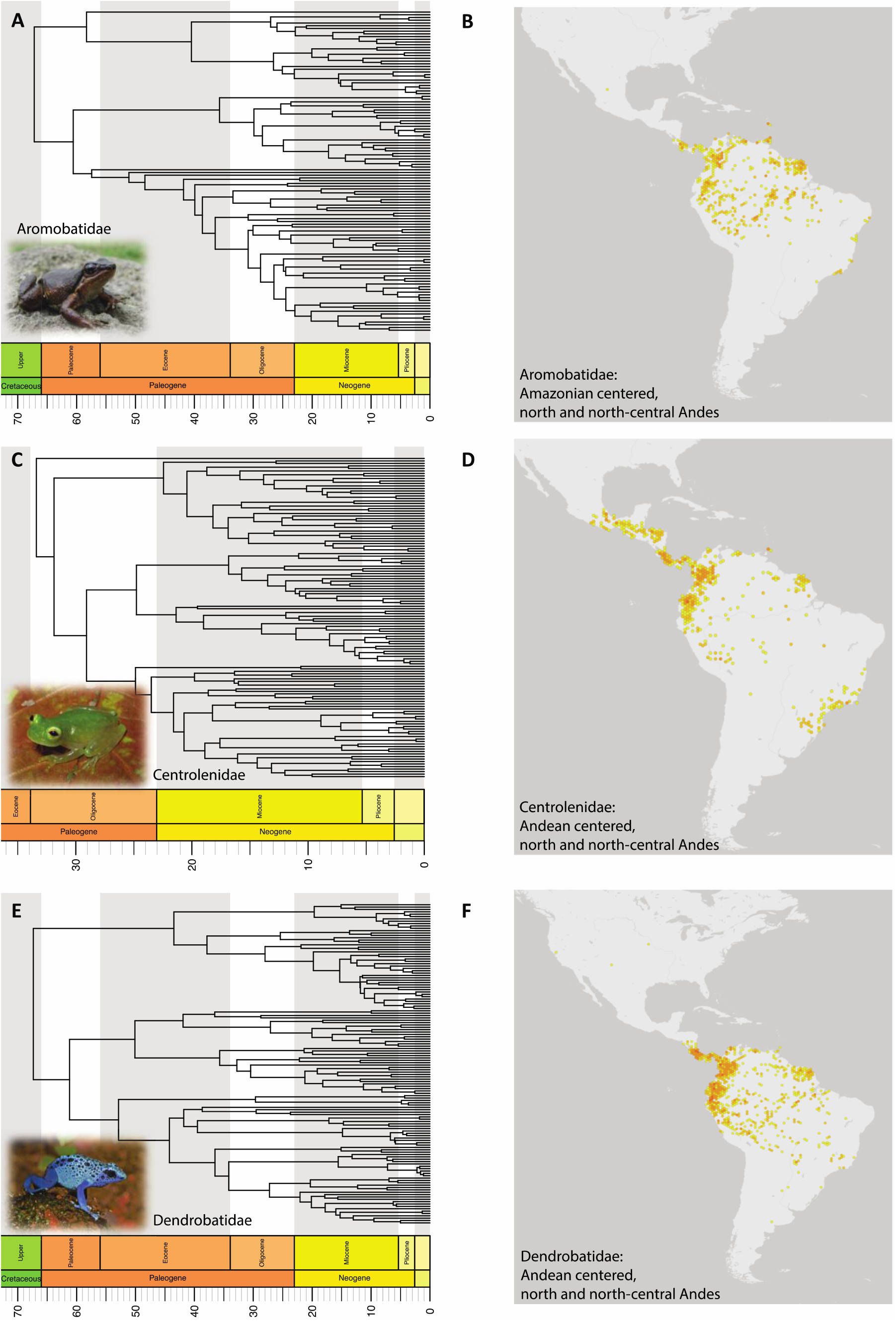
Distribution ranges of Aromobatidae, Centrolenidae and Dendrobatidae. From GBIF (the Global Biodiversity Information Facility, www.gbif.com).

**Supplementary Figure 2.**
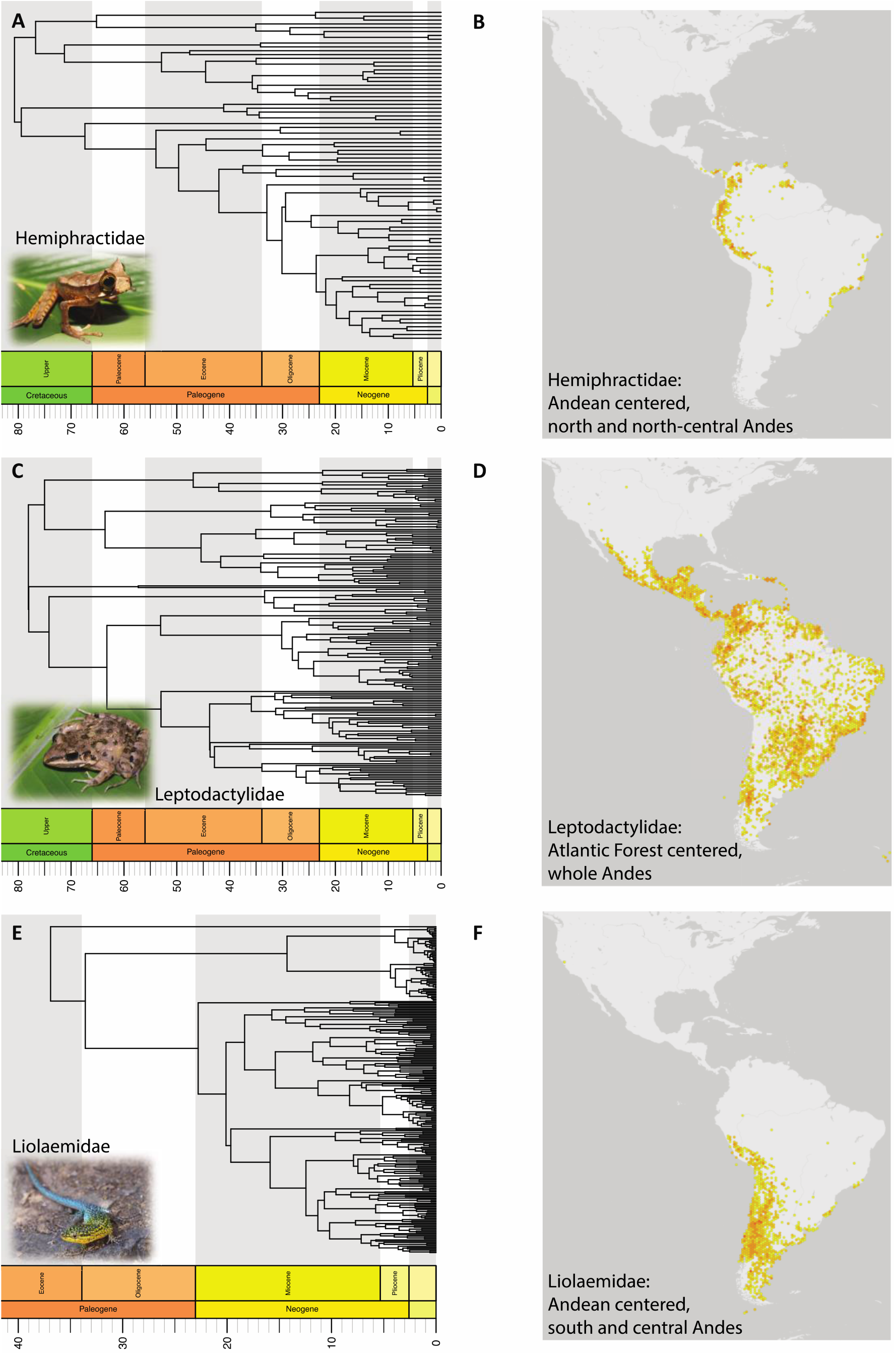
Distribution ranges of Hemiphractidae, Leptodactylidae, Liolaemidae. From GBIF (the Global Biodiversity Information Facility, www.gbif.com).

